# Genetic evidence that uptake of the fluorescent analog 2NBDG occurs independently of known glucose transporters

**DOI:** 10.1101/2021.12.13.472409

**Authors:** Lucas J. D’Souza, Stephen H. Wright, Deepta Bhattacharya

## Abstract

The fluorescent derivative of glucose, 2-Deoxy-2-[(7-nitro-2,1,3-benzoxadiazol-4-yl)-amino]-D-glucose (2NBDG), is a widely used surrogate reagent to visualize glucose uptake in live cells at single cell resolution. Using a model of CRISPR-Cas9 gene editing in 5TGM1 myeloma cells, we demonstrate that ablation of the glucose transporter gene *Slc2a1* abrogates radioactive glucose uptake but has no effect on the magnitude or kinetics of 2NBDG import. Extracellular 2NBDG, but not NBD-fructose was transported by plasma cells into the cytoplasm suggesting specific activity that is unlinked to glucose import and that of chemically similar compounds. RNA-Seq analysis of primary plasma cells and the 5TGM1 myeloma cell line revealed expression of other candidate glucose transporters. Yet, deletion of these transporters individually or in combination with one another also had no impact on 2NBDG uptake. Ablation of the genes in the *Slc29* and *Slc35* families of nucleoside and nucleoside sugar transporters as well as the ATP-binding cassette (ABC) transporter family also failed to impact 2NBDG import. Thus, cellular uptake of 2NBDG is promoted by an unknown mechanism and is not a faithful indicator of glucose transport.

## Introduction

Glucose is a critical nutrient for fueling cellular metabolism. After its import, glucose can be catabolized through a multitude of metabolic pathways. For example, glucose can be broken down by glycolysis into pyruvate that in turn is oxidized to fuel the tricarboxylic acid (TCA) cycle and generate ATP (1). Glucose is also important for generating intermediate sugars which serve as raw materials for the synthesis of nucleotides and fatty acids as well as glycosylation of cellular proteins (2). Glucose uptake and catabolism are key defining features of cells, distinguishing between developmental stages of lineages, tumors versus normal tissues, and activation status.

Definitive methods to measure glucose import have classically involved the use of isotope-labelled derivatives of glucose (3–5). A shortcoming to these compounds, however, is their rapid breakdown or export from the cell, and an inability to distinguish glucose uptake at a single cell level. Fluorescent derivatives of glucose, which allows for visualization and cytometric estimation of glucose uptake in cells, can potentially overcome these problems (6). The most popular of these compounds is 2-deoxy-2-(7-Nitro-2,1,3-benzoxadiazol-4-yl)amino-D-glucose (2NBDG), in which the 2-hydroxyl group of D-glucose is replaced with a fluorescent 7-Nitrobenzofurazan group (7). This compound was first characterized in *E. coli* where it competed with D-glucose for import via a mannose or a glucose/mannose transporter system (7,8). Since its discovery, it has been used across various mammalian cell types and *in vitro* culture models as a surrogate for glucose uptake. In plasma cells, we used this analog to demonstrate that 2NBDG+ long-lived plasma cells showed elevated spare respiratory capacity relative to their 2NBDG-short-lived counterparts, thereby linking glucose uptake with plasma cell longevity (9,10). Further, as compared to other markers of murine plasma cells, 2NBDG positivity correlated well with the longevity of the plasma cell subset (11). Yet despite its widespread use, the specificity has not been definitively shown for 2NBDG uptake through glucose transporters.

Glucose import into eukaryotic cells can take place via three families of transporters: first, the sodium-glucose linked symporters of the SGLT/SLC5 family of transporters; second, the newly characterized SWEET family of glucose transporters of the SLC50 family; and lastly, the well characterized GLUT/SLC2 family of sugar transporters (12,13). Through much of B cell development and activation, glucose uptake is mediated by the *Slc2* family member GLUT1, which is thus likely to be the chief glucose transporter in plasma cells (14). Our assumption had been that GLUT1 would be the likeliest candidate transporter for 2NBDG uptake as well. Contrary to our expectations, we demonstrate in this study that disruption of GLUT1 expression led to loss of glucose import but had no effect on 2NBDG uptake. Ablation of other candidate transporters also did not affect 2NBDG uptake. These data are consistent with two recent pharmacological and knockdown studies (15,16). We conclude that 2NBDG is actively transported into cells independently of known glucose transporters. Therefore, 2NBDG should not be used as a proxy for glucose uptake by mammalian cells.

## Materials and methods

### Ethics statement

All animal procedures carried out in this manuscript were approved and carried out based on guidelines provided by the Institutional Animal Care and Use committee at The University of Arizona (approval 17-266). Euthanasia was performed by administering carbon dioxide at a rate of 1.5L/minute in a 7L chamber until 1 minute after respiration ceased. Mice were then cervically dislocated to ensure death.

### Mice

C57BL/6N mice were purchased from the Charles River laboratories and housed under specific pathogen free conditions. Experiments were carried out on sex-matched mice between 8-12 weeks of age. For experiments involving *in vivo* NBD-metabolite uptake, mice were injected intravenously with 100μg of either 2NBDG or 1NBDF (both from Cayman chemical) and euthanized after 20 minutes.

### Cell lines and culture

The mouse myeloma line 5TGM1 was a gift from Michael H Tomasson at the Washington University in St. Louis (17). Cells were cultured in RPMI (Gibco) containing 10% fetal bovine serum (PEAK Serum), 2mM L-alanyl-L-glutamine, 1mM sodium pyruvate, minimal non-essential amino acids, penicillin, and streptomycin. Cells were maintained in a T-25 flask at 37°C with 5% CO_2_ and split every 4 days at a ratio of 1:10. Cas9-expressing 5TGM1 cells were generated by spin infecting 2×10^6^ 5TGM1 cells with lentiCas9-BLAST lentivirus and 8μg/mL of Polybrene (Millipore Sigma) at 2500rpm for 90 minutes followed by selection in 10μg/mL Blasticidin-S-HCl (Gibco). Guide RNA (gRNA) containing lentiviruses were introduced into these cells by similar spin infections, followed by selection with 10μg/mL Puromycin (Gibco) at 48 hours. Alternatively, 5TGM1 cells were spin-infected with lentiCRISPRv2-mCherry lentivirus to generate an Cas9-expressing line that showed mCherry fluorescence. These cells were purified from uninfected cells by fluorescence-activated cell sorting. For assays involving 2NBDG uptake, 1×10^6^ 5TGM1 cells were cultured with 20μg/mL 2NBDG (Cayman Chemical) in complete media for 1 hour at 37°C followed by antibody staining. LentiX-293T cells (632180, Takara Bio) were used for lentivirus packaging and assembly. They were maintained in DMEM containing 10% fetal bovine serum, L-alanyl-L-glutamine, minimal non-essential amino acids, sodium pyruvate, penicillin, and streptomycin. Cells were cultured in a 100mm petri dish and split using 0.05% Trypsin-EDTA (Millipore Sigma) when it exhibited greater than 80% confluence.

### Plasmids and lentivirus generation

lentiCas9-Blast, lentiCRISPRv2-mCherry, and lentiGuide-puro were all used in this study (52962, 52961, and 52963 respectively; Addgene). gRNA sequences were chosen from the existing mouse Brie library or designed using the CRISPick platform and cloned into lentiGuide-puro as described previously (18–20). In brief, 20-mer gRNA sequences were introduced into primers (Millipore Sigma) and after hybridization were cloned into Esp3I-digested lentiGuide-puro using the NEBuilder® HiFi DNA Assembly Master mix (E2621, NEB). Successful constructs were verified in plasmid from isolated colonies by sequencing with the LKO.1 5’ primer. Constructs used for generation of the *Slc2a3, Slc2a5, Slc2a6,* and *Slc2a8* knockout cell line were prepared by digesting lentiGuide-puro constructs with already cloned gRNAs sequences with BsiWI-HF and MluI-HF (R0553 and R3198, NEB) to exclude the puromycin N-acetyltransferase gene. Sequences encoding fluorescent proteins were then cloned into these digested vectors to generate *Slc2a3*-lentiGuide-dsRed, *Slc2a5*-lentiGuide-mMaroon, *Slc2a6*-lentiGuide-mCherry, and *Slc2a8*-lentiGuide-tagBFP.

LentiX-293T cells (632180, Takara Bio) were cultured to 60% confluence in 10cm^2^ dishes and transfected using 30μL GeneJuice Transfection reagent (Millipore Sigma), 1.75μg of pMD2.G (12259, Addgene), 3.25μg of psPAX2 (12260, Addgene), and 5μg of the lentiviral construct as per the manufacturer’s instructions. Media was changed at 6 hours post transfection and supernatants collected at 48 and 72 hours and filtered through a 0.45μ syringe. Filtered supernatants were then mixed in a 5:1 ratio with 25% polyethylene glycol-8000 (Millipore Sigma) in 1×PBS and incubated overnight at 4°C. Samples were then spun down at 2500rpm for 20 minutes and pellets resuspended in 100μL 1xPBS. Aliquots were then frozen at −80°C until use.

### Flow cytometry

All fluorescence associated cell sorting was carried out on a BD FACS Aria II and analysis on a BD LSR II (Becton Dickinson). Single cell suspensions from spleens and bone marrows of mice were prepared and erythrocytes lysed with an ammonium chloride-potassium (ACK) lysis buffer. Lymphocytes were then isolated using a Histopaque-1119 (Millipore Sigma) and cells were suspended in 1x PBS containing 5% adult bovine serum (FACS buffer). Cultured 5TGM1 cells were washed once in FACS buffer prior to antibody staining. The following anti-mouse antibodies were used: CD138-Phycoerythrin (PE), -Allophycocyanin (APC), or -BV510 (281-2, Biolegend) and B220-BV421 (RA3-6B2, Biolegend). When staining cultured cells, propidium iodide (Millipore Sigma) or Zombie UV (Biolegend) were used to exclude dead cells from the analysis. To stain for GLUT1, cells were first fixed in 2% paraformaldehyde (Electron Microscopy Services) and then stained with an unconjugated GLUT1 monoclonal antibody (SPM498, Thermo Fisher Scientific) followed by a Rat anti-mouse IgG2a -Alexa Fluor 647 detection antibody (SB84a, Southern Biotech). Cells stained with the detection antibody alone were used as an isotype control. Data was analyzed using the FlowJo software (Becton Dickinson).

### Imagestream analysis

2NBDG treated 5TGM1 cells expressing mCherry with control or *Slc2a1*-targeting gRNA were stained for surface CD138. Samples were then analyzed on an Imagestream^X^ Mk II (Luminex) and 3000 events recorded per group at 60x magnification. Raw information files analyzed on the IDEAS v.6.3 software (Luminex) and similarity morphology indices calculated using the nuclear localization wizard. Mean similarity morphology indices for groups across three experiments were then graphed using Prism 9.2 (Graphpad).

### ^14^C-Glucose uptake assays

1×10^6^ 5TGM1 cells were resuspended in glucose-free incomplete RPMI (Gibco) in a microcentrifuge tube and incubated briefly in a water bath set at 37°C. ^14^C Glucose (Perkin Elmer) was introduced into these cultures to reach a final concentration of 0.5 μCi/mL and cells were allowed to incubate with shaking for 30 minutes. Cells were then washed once with 1x PBS (HyClone) and then lysed in a solution containing 0.5N NaOH and 1% SDS for 30 minutes at room temperature. After neutralization with 1N HCl, cell lysate was loaded onto a 96-well plate (Grenier-BioOne) and mixed with MICROSCINT^TM^-20 scintillation fluid (Perkin Elmer). Samples were then incubated at room temperature for 2 hours and incorporated radioactivity analyzed on a 1450 MicroBeta TriLux Microplate Scintillation and Luminescence counter (Perkin Elmer).

### Next generation sequencing

Genomic DNA was isolated from gRNA-transduced cultures at the time of assay using a DNA extraction kit (IBI Scientific). Primers were designed +/-150bp from the PAM site of the targeting gRNA and used to amplify 300bp amplicons from 1μg genomic DNA using the Q5® DNA polymerase (M0491, NEB) for 25 cycles. Amplicons were then gel extracted, amplified with the same primers and reaction conditions for 2 cycles, and purified by gel extraction. Equimolar concentrations of amplicons from different reactions were then pooled and 300fmol of the sample was end-prep treated and ligated with barcodes and adapter sequences as per the ligation sequencing kit protocol (SQK-LSK109, Oxford Nanopore). For some libraries, barcodes were introduced into the samples using a modified version of the ligation sequencing kit protocol (NBD-104, Oxford Nanopore). Samples were then loaded onto primed SpotON flow cells and sequenced on a MinION Mk1c (Oxford Nanopore) with a read filter of 250-500bp and high-accuracy basecalling. Reads in resultant FASTQ files were mapped to the region of interest and indels calculated using the CRISPResso2 analysis pipeline (21). Unmodified, in-frame, and frame-shift mutation containing read frequencies were graphed as stacked columns using Prism 9.2 (Graphpad).

### Statistical analysis

All statistical analysis was carried out on Prism 9.2 (Graphpad). Specific tests used and significance is indicated in the figures and accompanying figure legends.

## Results

### GLUT1 does not mediate uptake of 2NBDG

The glucose transporter GLUT1, encoded by the gene *Slc2a1,* is highly expressed in various models of multiple myeloma, a transformed counterpart of long-lived plasma cells (22). To test if GLUT1 is responsible for glucose uptake, we devised a model of CRISPR-Cas9 mediated gene deletion in the 5TGM1 mouse myeloma system. Using lentiviral transduction, we engineered these cells to constitutively express the Cas9 protein, and after selection, introduced guide RNAs (gRNA) targeting genes of interest through lentiviral transduction. Accordingly, we introduced four different gRNAs that targeted exons 3, 4, and 5 of the *Slc2a1* locus into these cells. The observed frequency of GLUT1-positive cells in these cultures dropped to approximately 50% relative to that of a control gRNA transduced culture (Fig 1 A). This loss in expression of GLUT1 was accompanied by a significant decrease in ^14^C-glucose uptake (Fig. 1 B). Unexpectedly, the import of 2NBDG was unaffected in *Slc2a1*-targeted gRNA cultures (Fig 1 C). Further, we saw no change in the kinetics of 2NBDG uptake in these cells at any of the time points measured (Fig 1 D). These findings suggest that mechanisms or transporters other than GLUT1 can mediate uptake of 2NBDG.

**Figure 1:**
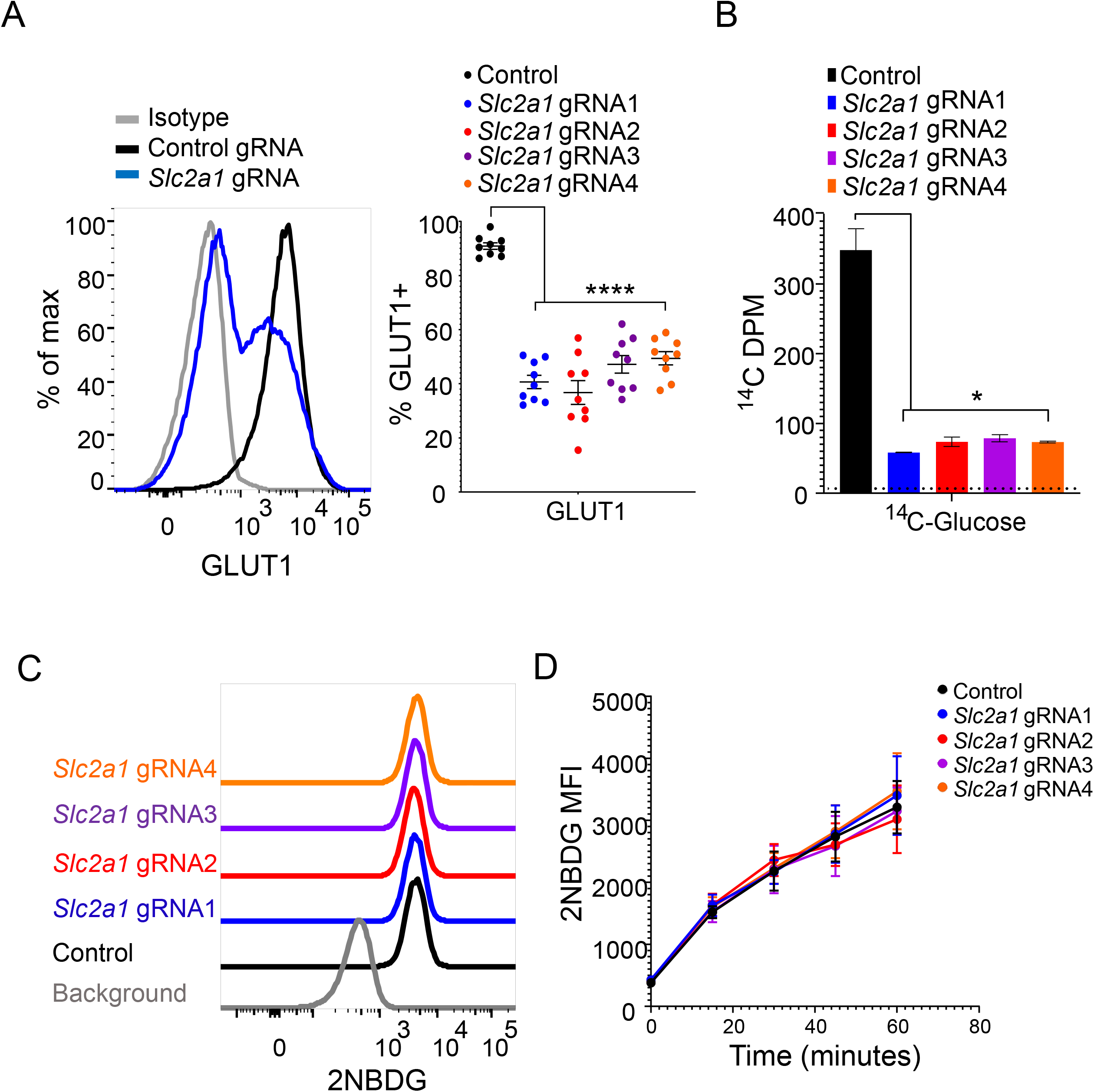
2NBDG uptake is unaffected by *Slc2a1* deletion in 5TGM1 cells. (A) GLUT1 staining of gRNA transduced cells. Representative histogram (left) showing GLUT1 expression on 5TGM1 cells transduced with a control gRNA (black) and a *Slc2a1* targeting gRNA (blue). Quantification of percent GLUT1 positive cells (right) in control and *Slc2a1* gRNA transduced cells across 9 independent experiments. Each circle represents a single group from a single experiment. ***p<0.0001 by Brown-Forsyth and Welch one-way ANOVA multiple comparisons test. (B) ^14^C-glucose uptake in gRNA transduced cells. Lysates from cells transduced with gRNAs as in (A) were cultured in ^14^C-glucose containing media for 30 minutes and estimated for ^14^C. Background counts indicated by the dotted line. Representative graph of three independent experiments. *p<0.05 by Brown-Forsyth and Welch one-way ANOVA multiple comparisons test. (C) Flow cytometric analysis of 2NBDG uptake in control gRNA (black) and *Slc2a1* gRNA (colored) transduced 5TGM1-Cas9 cells. Cells were cultured in media containing the metabolite for 60 minutes. Histogram representative of three independent experiments. (D) 2NBDG uptake in control gRNA (black) and *Slc2a1* gRNA (colored) transduced 5TGM1-Cas9 cells. Mean fluorescence intensity (MFI) +/− SEM shown for 0-, 15-, 30-, 45-, and 60-minutes post culturing with 2NBDG. Pooled data from three independent experiments. No significant differences observed with ordinary two-way ANOVA with Dunnett’s multiple comparison test.

### Plasma cells take up 2NBDG, but not 1-NBD-Fructose

We next considered the possibility that 2NBDG, which is hydrophilic, is endocytosed by cells in a non-specific manner rather than delivered to the cytoplasm via a transporter. To test this, we examined 5TGM1 cells treated with 2NBDG using imaging flow cytometry. We observed that 2NBDG was distributed evenly across the cytosol of cells and absence of the sugar transporter GLUT1 did not affect this distribution (Fig. 2 A). Further, 2NBDG showed high similarity morphology scores with a cytosolic mCherry versus the surface marker CD138, indicating that 2NBDG is cytosolic and likely imported via a transporter (Fig. 2 A). To test the specificity of this unknown transporter, we injected mice with 2NBDG or 1-NBD-fructose (1NBDF), a derivative of fructose in which the NBD moiety is attached to the C-1 sugar of fructose (Supplemental Figure 1, and 23). We observed 2NBDG uptake in both splenic and bone marrow plasma cells, with the latter showing a higher frequency of 2NBDG-positive cells (Fig. 2 B), as we previously reported (9,10). Plasma cells, however, did not detectably import 1NBDF (Fig. 2 B). These data suggest that plasma cells import 2NBDG specifically, though this does not appear to be mediated by GLUT1.

**Figure 2:**
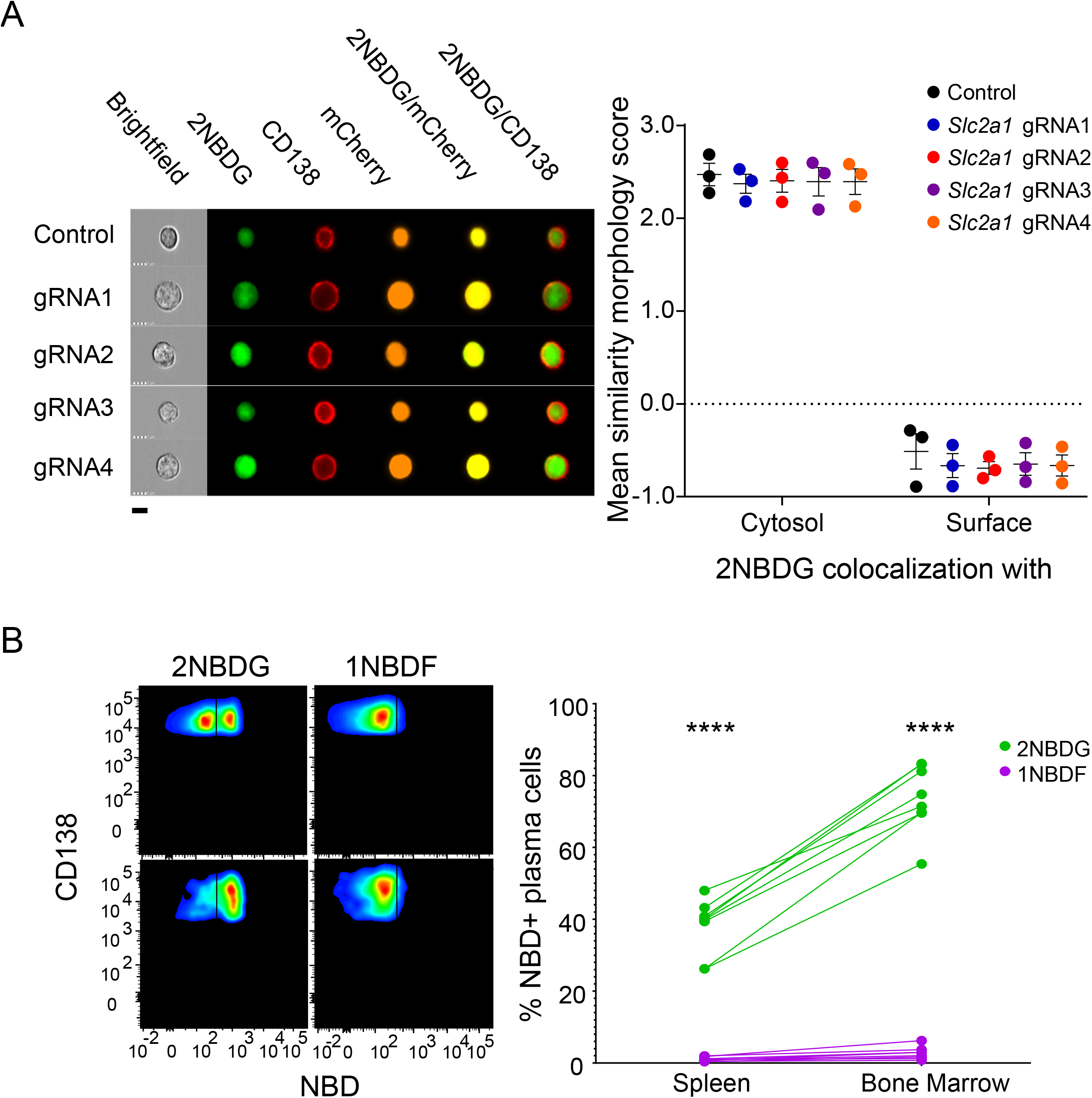
Plasma cells specifically take up 2NBDG and retain it in the cytosol. (A) 2NBDG is retained in the cytosol of cells. 5TGM1 cells previously transduced with a Cas9-T2A-mCherry lentivirus followed by control gRNA or *Slc2a1* gRNA lentiviruses were treated with 2NBDG, stained for surface CD138, and analyzed on the Imagestream. Representative images of each gRNA group (left) at 60x magnification are shown with overlays for 2NBDG/CD138 and 2NBDG/mCherry. The black bar indicates a length of 7 microns. Mean similarity morphology indices were quantified and represented (right). Each dot indicates the mean value for a group in a single experiment. No significant differences observed with ordinary two-way ANOVA with Dunnett’s multiple comparison test. (B) Mice were injected with 100μg of either 2NBDG or 1NBDF and assessed for NBD fluorescence in splenic and bone marrow plasma cells. Representative flow cytometry plots (left) on splenic (top row) and bone marrow (bottom row) CD138+ plasma cells showing gated percent NBD-positive cells. Quantification of NBD-positive plasma cell percentages (left) in the spleen and bone marrow and groups from each mouse are connected by a line. Data from three independent experiments with n=8 mice in both groups. ****p<0.0001 by ordinary two-way ANOVA with *post hoc* Sidak’s multiple comparison test.

### 2NBDG uptake does not depend on other glucose transporters

We next hypothesized that 2NBDG uptake into cells could take place via other transporters involved in glucose uptake. Using RNA-seq data, we identified known and putative glucose transporters of the SLC2, SLC5, and SLC50 families that are expressed by 5TGM1 and/or primary plasma cells (9,10,24). Of the 13 members of the *Slc2* family in mice, primary plasma cells and/or myeloma cells expressed SLC2A3, SLC2A5, SLC2A6, and SLC2A8 as candidate glucose transporters in addition to SLC2A1 (Fig. 3 A). Expression of SLC5 family members was not detected. We also observed high levels of expression of SLC50A1, though this transporter is primarily involved in glucose efflux rather than import (13).

**Figure 3:**
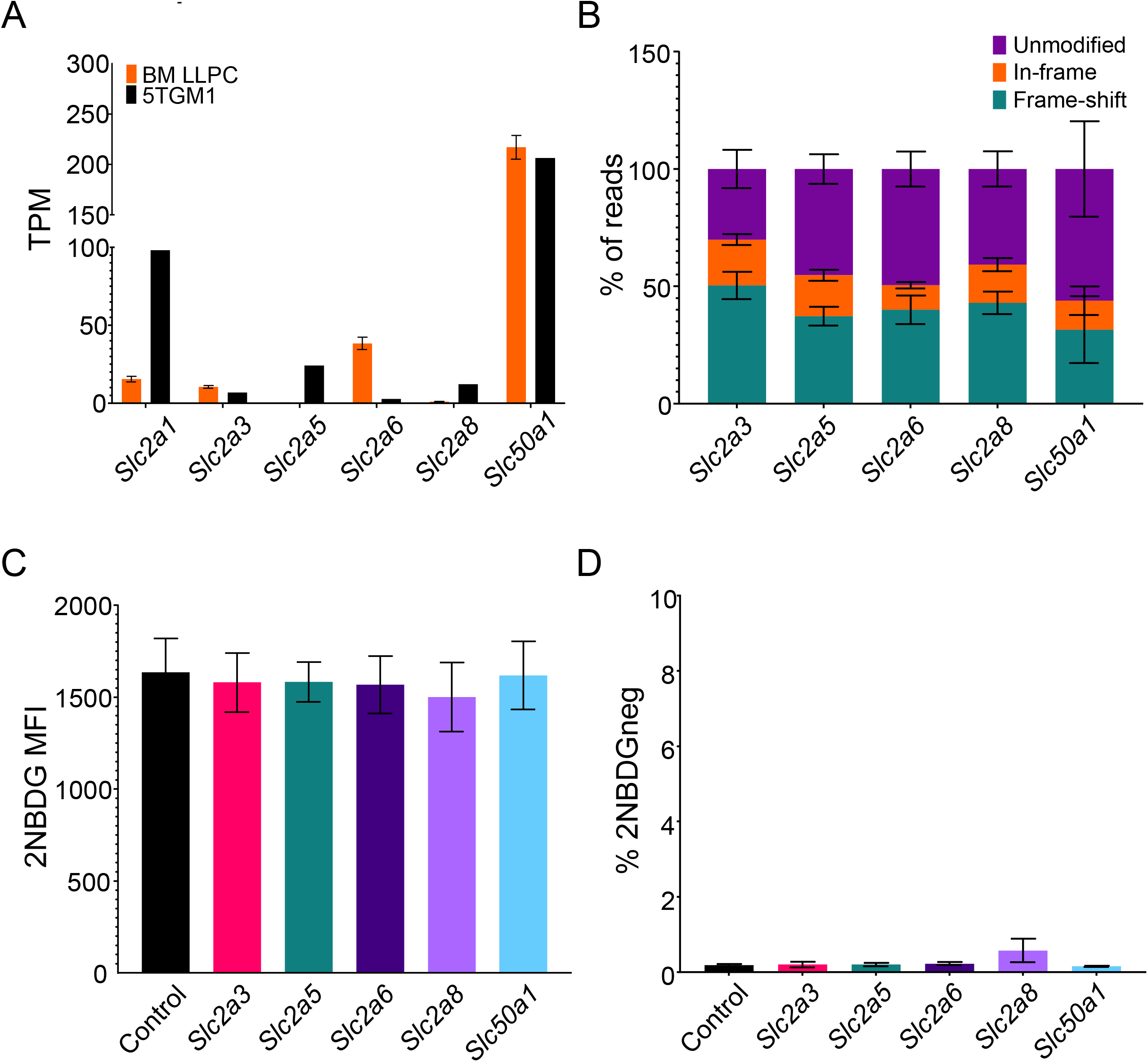
2NBDG uptake takes place independently of sugar transporters. (A) TPM values of sugar transporters in *ex vivo* bone marrow plasma cells (orange) and 5TGM1 cells (black). Mean values +/− SEM are shown. (B) Quantification of gene modifications in gRNA transduced cultures. Exons of indicated genes were PCR amplified and sequenced to assay for in-frame and frame shift mutations. Mean values +/− SEM shown for each of the genes and modifications within it. Pooled data from three experiments. (C) 2NBDG uptake in sugar transporter deleted cultures. 5TGM1-Cas9 cells were transduced with control gRNA (black) or gRNAs targeting *Slc2a3* (pink), *Slc2a5* (blue), *Slc2a6* (violet), *Slc2a8* (lilac), and *Slc50a1* (cyan). MFIs across three independent experiments are quantified and displayed with SEM. No significant differences observed with Brown-Forsyth and Welch one-way ANOVA multiple comparisons test. (D) Frequency of 2NBDG-negative cells for groups in (C). 2NBDG-gate drawn based on control cells cultured devoid 2NBDG in complete media. Mean values +/− SEM are shown. No significant differences observed with Brown-Forsyth and Welch one-way ANOVA multiple comparisons test.

To test out if these transporters are responsible for 2NBDG import, we transduced gRNAs targeting these genes into Cas9-expressing 5TGM1 cells and scored for 2NBDG uptake by flow cytometry. Sequencing of gRNA targets in these cultures demonstrated 31-50% frameshift mutations, confirming efficient ablation of the intended transporters (Fig. 3 B). Disruption of these genes, however, did not affect 2NBDG uptake (Fig. 3 C). Moreover, the frequency of 2NBDG-negative cells in all cultures were similar to those seen in control gRNA transduced cultures (Fig. 3 D).

A possible explanation for the unaltered 2NBDG uptake in the *Slc2a1, Slc2a3, Slc2a5, Slc2a6,* and *Slc2a8* deleted cultures is functional redundancy between these sugar transporters. As a result, loss of one of these genes might be compensated by activity of another transporter(s). To test this possibility, we generated 5TGM1 cells carrying gRNAs targeting all the expressed members of the SLC2 family. Because SLC2A1-deficient cells show reduced viability, we generated a cell line that was ablated for *Slc2a3, Slc2a5, Slc2a6,* and *Slc2a8* first using lentiviral transduction followed by FACS of the reporter positive cells. After sorting and expansion of this line, we then introduced lentiviruses expressing a gRNA targeting *Slc2a1* and examined for 2NBDG uptake in these cells 4 days after transduction. Analysis of the number of reads in these cultures showed that *Slc2a1* sequences had an average of 50% frameshift mutations, while the other *Slc2* members had 54-65% frameshifts in their respective sequences (Fig. 4 A). 2NBDG uptake was equivalent in these cells relative to controls (Fig. 4 B-D). Thus, known glucose transporters are not required for 2NBDG uptake.

**Figure 4:**
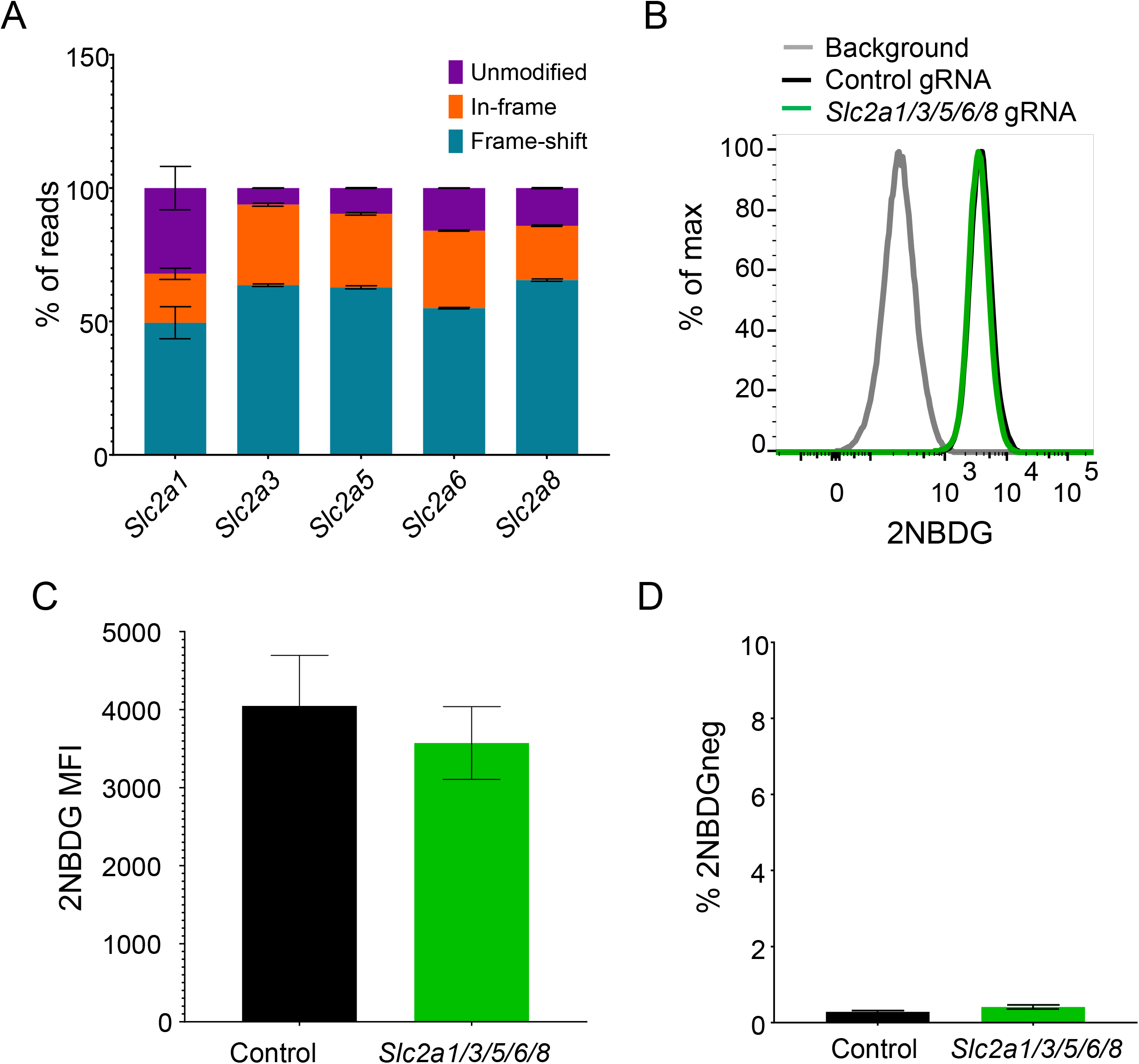
Combined deletion of sugar transporters does not affect 2NBDG uptake. (A) Quantification of in-frame and frame shift mutations in *Slc2a1, Slc2a3, Slc2a5, Slc2a6,* and *Slc2a8* genes of the *Slc2a1/3/5/6/8* deleted cultures. Pooled data from three experiments. (B) 2NBDG uptake in *Slc2a1/3/5/6/8* cultures. Representative histogram (left) showing control gRNA (black) and the five gRNAs transduced culture (green). Pooled MFI +/− SEM for both groups across three independent experiments are depicted (right). No significance observed with the Mann-Whitney non-parametric t test. (C) Frequencies of 2NBDG-negative cells in cultures described in (A). 2NBDG-gating carried out as in Figure 3 (D). Mean values +/− SEM are shown for both groups. No significance observed with the Mann-Whitney non-parametric t test.

### Nucleoside and nucleoside-sugar transporters do not transport 2NBDG

Given the utility of 2NBDG import as a marker of plasma cell longevity, we performed experiments to test other candidate transporters. We first reasoned that the structure of 2NBDG mimicked a nucleotide or nucleotide sugar and likely was imported though their transporters which are part of the *Slc29* and *Slc35* families (Supplementary Figure 1). RNA-seq data showed *Slc29a1* and *Slc29a3* expression in both primary bone marrow plasma cells and 5TGM1 cells (Fig. 5 A). Moreover, 17 of the 27 known *Slc35* members in mice were also expressed (Fig. 5 A). CRISPR-Cas9 ablation of these transporters led to 20-59% frameshift mutations in sequences from 5TGM1 cultures (Fig. 5 B). However, none of these mutations prevented 2NBDG uptake (Fig. 5 C-D). Thus, the *Slc29* or *Slc35* families of nucleotide and nucleotide sugar transporters are not required for 2NBDG uptake.

**Figure 5:**
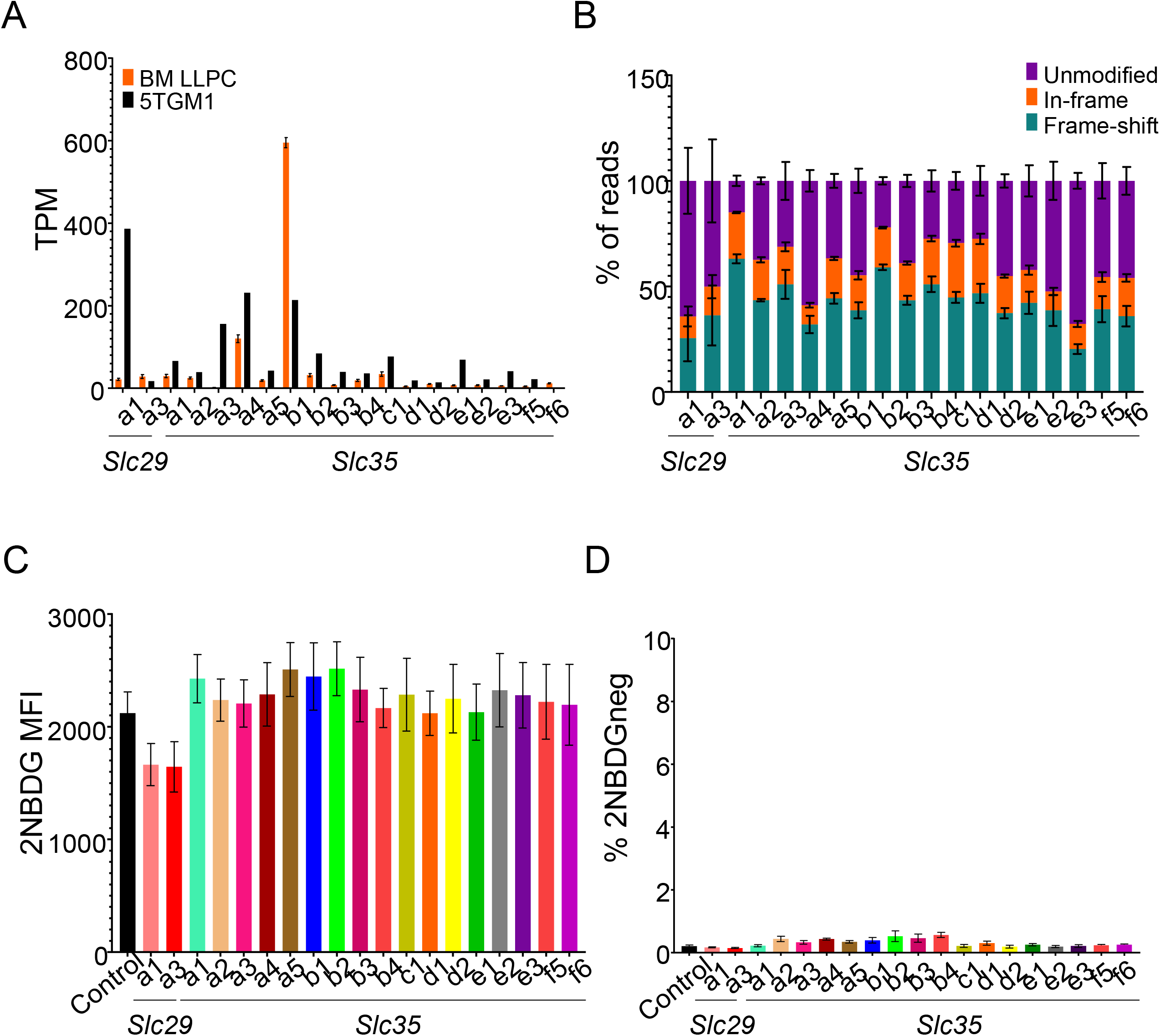
2NBDG is not imported through nucleoside or nucleoside-sugar transporters. (A) TPM values of nucleoside and nucleoside-sugar transporters in *ex vivo* bone marrow (orange) plasma cells and 5TGM1 cells (black). Mean values +/− SEM are shown. (B) Estimation of indels in nucleoside and nucleoside-sugar transporter deleted cultures done as in Figure 3 (B). Mean values +/− SEM shown for each of the genes and modifications within it. Pooled data from three independent experiments. (C) 2NBDG uptake in deleted cultures. 5TGM1-Cas9 cells transduced with gRNAs targeting indicated nucleoside or nucleoside-sugar transporters (colored) or control gRNA (black). 2NBDG MFIs across three independent experiments are quantified and displayed with SEM. No significant differences observed with Brown-Forsyth and Welch one-way ANOVA multiple comparisons test. (D) Frequencies of 2NBDG negative cells in cultures described in (C). Pooled data from three experiments. No statistical significance observed with the Mann-Whitney non-parametric t-test.

### 2NBDG is not imported by the ABC family of transporters

The broad variety of substrates transported via the ATP-binding cassette (ABC) family of transporters led us to next test if one of these transporters are involved in 2NBDG import (25,26). Of the 50 known transporters of this family in mice, we identified 10 that were expressed by both primary plasma cells and 5TGM1 cells (Fig. 6 A). We observed 24-62% frameshift mutations in the target gene sequences of gRNA transduced 5TGM1-Cas9 cultures (Fig. 6 B). Yet we observed no significant difference in 2NBDG uptake or the frequency of 2NBDG negative cells in ABC transporter-deleted cultures relative to control gRNA transduced cultures (Fig. 6 C, D). Together, our data demonstrate that 2NBDG uptake, while specific, is not mediated by known sugar, nucleotide, nucleotide sugar, or ABC transporters.

**Figure 6:**
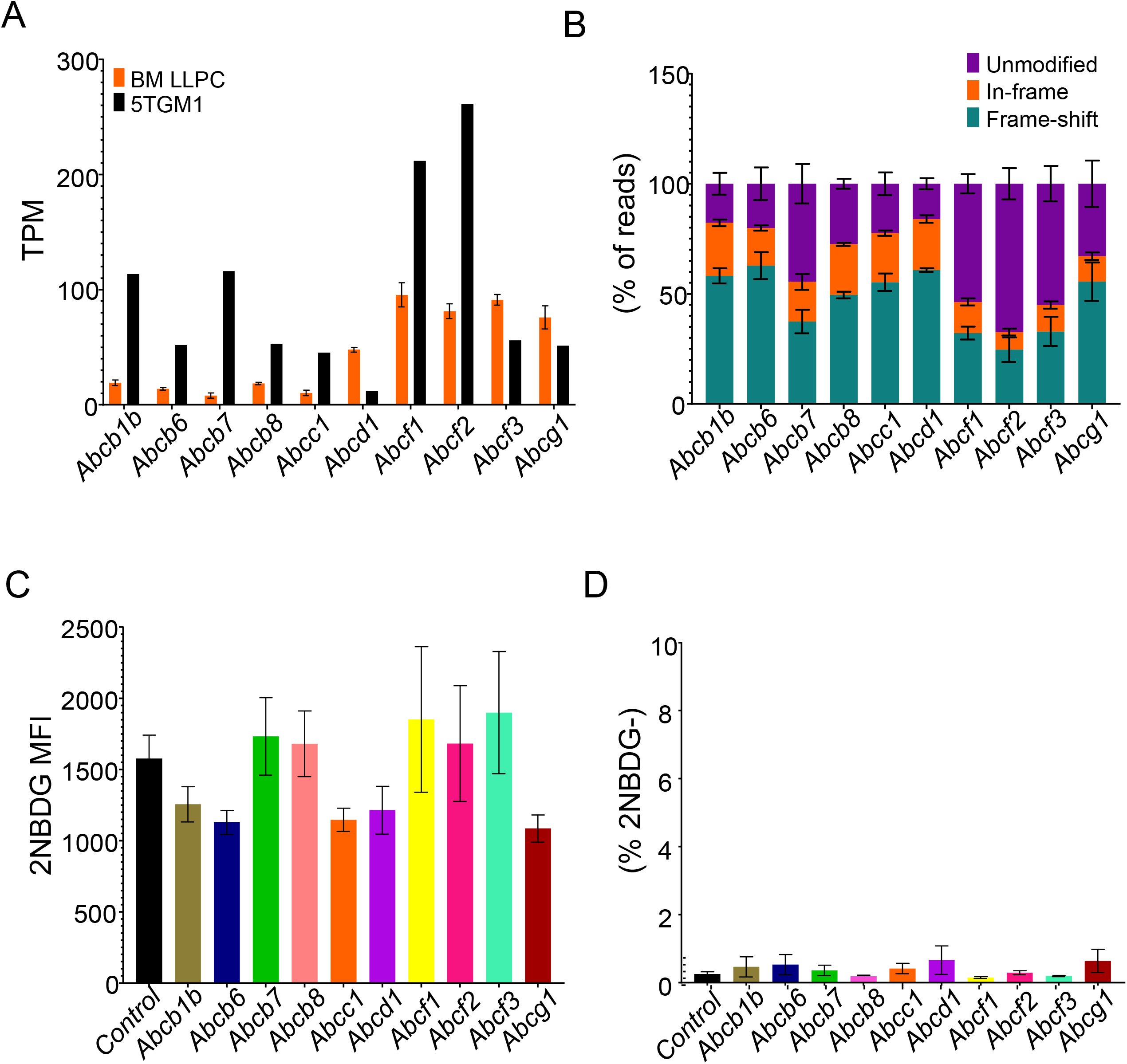
ABC transporters do not mediate uptake of 2NBDG in 5TGM1 cells. (A) TPM values of ABC transporters in *ex vivo* bone marrow plasma cells (orange) and 5TGM1 cells (black). Mean values with SEM are shown (B) Estimation of indels in ABC transporter deleted cultures done as in Figure 3 (B). Mean values +/− SEM shown for each of the genes and modifications within it. Pooled data from three independent experiments. (C) 2NBDG uptake in deleted cultures. 5TGM1-Cas9 cells transduced with gRNAs targeting indicated ABC transporters (colored) or a control gRNA (black). 2NBDG MFIs across three independent experiments are quantified and displayed with SEM. No significant differences observed with Brown-Forsyth and Welch one-way ANOVA multiple comparisons test. (D) Frequencies of 2NBDG negative cells in ABC transporter deleted cultures described in (C). Pooled data from three experiments. No statistical significance observed with the Mann-Whitney non-parametric t-test.

## Discussion

The use of fluorescent derivatives of glucose like 2NBDG has been a valuable tool in visualizing glucose uptake in cells and *in vivo.* While direct evidence for its uptake through glucose transporters exists in bacterial systems, a formal confirmation of its import through mammalian sugar transporters has not been demonstrated genetically (8). Using a model of CRISPR-Cas9 in myeloma cells, we show that the sugar transporter GLUT1 is important for glucose uptake but has no role in 2NBDG transport in these cells. As glucose can be imported into cells via other sugar transporters, we test their role in 2NBDG uptake and find no difference in 2NBDG intensity between control and deleted cultures. As such, our findings show a disconnect between glucose uptake and 2NBDG transport in mammalian cells and strongly advise against any correlation between these two distinct cellular processes.

During the course of this work, two reports were published by independent groups that arrived at similar conclusions. The first report by Sinclair *et al.* demonstrated that double-positive (DP) thymocytes did not take up any glucose as compared to activated CD8+ T cells in culture (15). The 2NBDG uptake by cells in these cultures, however, showed the exact opposite trend and was unaffected by pharmacological inhibition of glucose transporters (15). The second group took advantage of the L929 fibroblast line, that expresses GLUT1 as its sole glucose transporter (27). Using pharmacological inhibitors, siRNA mediated knockdowns, and GLUT1 overexpression, the authors were able to modulate glucose uptake in these cells, but not of 2NBDG or another fluorescent derivative of glucose, 6NBDG (16). Our findings provide an independent genetic confirmation of both reports, and further extend the disparity between glucose and 2NBDG uptake to other glucose transporters within the *Slc2* and *Slc50* families. The use of CRISPR-Cas9 to induce gene deletions in our system offers a robust means to provide this independent confirmation.

In plasma cells, 2NBDG uptake marks longer lived subsets and identifying its transporter could provide insights into mechanisms regulating the longevity of these cells. Long-lived plasma cells do possess more glucose-dependent spare respiratory capacity than do short-lived plasma cells (10). However, our original conclusions that long-lived plasma cells import more glucose than their short-lived counterparts will need to be revisited. The as-yet unidentified transporter seems specific, as our data indicate that cells deposit 2NBDG in the cytosol. We attempted an unbiased genome-wide gRNA library screen in the 5TGM1 cell line but were unable to identify any known candidates in sorted 2NBDG-negative cultures (data not shown). The lack of any putative targets in the screen suggests that 2NBDG might be imported into cells via multiple functionally redundant transporters. Such transporters might be identified through an overexpression library screen in cells that intrinsically do not take up 2NBDG. As the mechanisms of 2NBDG uptake remain unclear, we strongly advise that, despite their convenience, these assays not be used as surrogates for glucose uptake.

## Acknowledgements

We wish to thank the flow cytometry core at the University of Arizona for their assistance with flow cytometry.

## Author contributions

Conceptualization: Lucas D’Souza, Deepta Bhattacharya

Data curation: Deepta Bhattacharya

Formal analysis: Deepta Bhattacharya

Funding acquisition: Stephen Wright, Deepta Bhattacharya

Investigation: Lucas D’Souza, Deepta Bhattacharya

Methodology: Lucas D’Souza

Supervision: Stephen Wright, Deepta Bhattacharya

Writing- original draft: Lucas D’Souza

Writing-review and editing: Lucas D’Souza, Stephen Wright, Deepta Bhattacharya

## Supporting Information

**Supplementary Figure 1:**
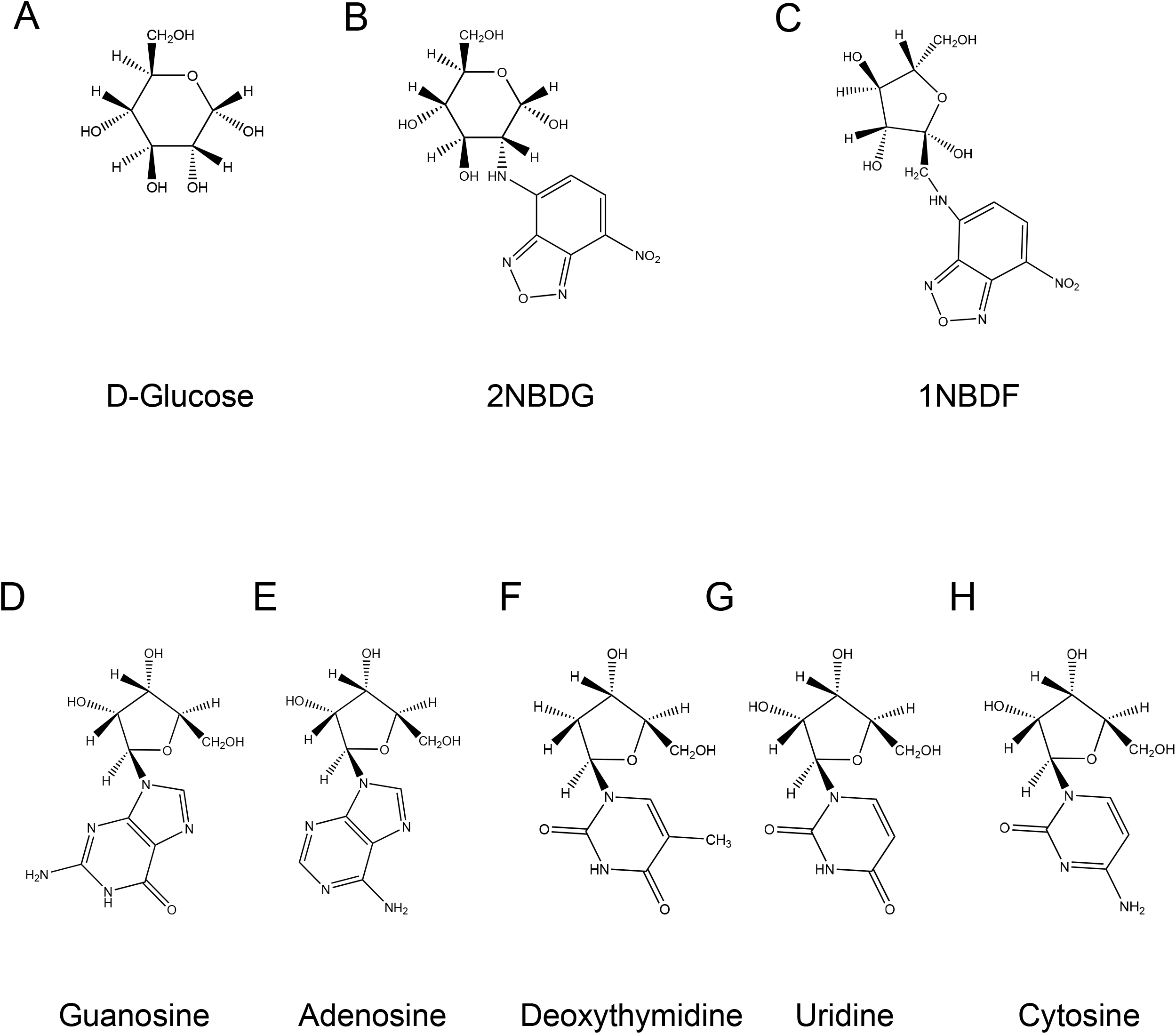
Structures of 2NBDG and 1NBDF in relation to glucose and naturally occurring nucleosides. Chemical structures of (A) D-Glucose, (B) 2NBDG, (C) 1NBDF, (D) Guanosine, (E) Adenosine, (F) Deoxythymidine, (G) Uridine, and (H) Cytosine. Structures generated using ChemDraw v.20.1.1

